# Soluble CTLA-4 raises the threshold for T-cell activation and modulates anti-tumour immunity

**DOI:** 10.1101/2023.06.05.543731

**Authors:** Paul T. Kennedy, Emma L. Saulters, Andrew D. Duckworth, Yeong Jer Lim, John F. Woolley, Joseph R. Slupsky, Mark S. Cragg, Frank J. Ward, Lekh N. Dahal

## Abstract

CTLA-4 is a crucial immune checkpoint receptor involved in the maintenance of immune homeostasis, tolerance, and tumour control. Antibodies targeting CTLA-4 have been promising treatment for numerous cancers, but the mechanistic basis of their anti-tumoral immune boosting effects are poorly understood. Although the *ctla4* gene also encodes an alternatively-spliced soluble variant (sCTLA-4), preclinical/clinical evaluation of anti-CTLA-4-based immunotherapies have not considered the contribution of this isoform. Here, we explore the functional properties of sCTLA-4 and evaluate the efficacy of isoform-specific anti-sCTLA-4 antibody targeting in murine cancer model. We show that expression of sCTLA-4 in tumour cells suppresses CD8^+^ T-cells *in vitro*, and accelerates growth and experimental metastasis of murine tumours *in vivo*. These effects were accompanied by modification of the immune infiltrate, notably restraining CD8^+^ T-cells in a non-effector state. sCTLA-4 blockade with isoform-specific antibody reversed this restraint, enhancing intratumoural CD8^+^ T-cell activation and cytolytic potential, correlating with therapeutic efficacy and tumour control. This previously unappreciated role of sCTLA-4 suggests that better understanding of the biology and function of multi-gene products of immune checkpoint receptors needs to be fully elucidated for improved cancer immunotherapy.

## Introduction

Cytotoxic T-lymphocyte–associated antigen 4 (CTLA-4) is a regulator of T-cell activation and was the first molecule successfully targeted for immune checkpoint therapy (1). However, the mechanisms by which it suppresses immune responses *in vivo* remain ill-defined (2,3). Constitutively expressed as a membrane protein on the surface of regulatory T-cells (T_reg_) and activated effector T-cells, CTLA-4 competes with CD28 for engagement of the B7 ligands CD80/CD86 on antigen presenting cells (APC). Such interaction reverses CD28-mediated co-stimulation of T cells via a mechanism involving induction of negative signal transduction (4), and is responsible for controlling autonomous activation of T-cells (5,6) as well cell-extrinsic regulation of distal T-cell populations (7,8). Thus, our current understanding of how CTLA-4 functions centres around its membrane-bound isoform structure (9). Here it is important to note that transcripts of the *ctla4* gene in humans and mice can be alternatively spliced to yield membrane-bound and secreted variants, the latter resulting from deletion of the exon encoding the transmembrane domain and a frameshift giving rise to a unique C-terminal sequence (10,11). Why this is important is because currently available anti-CTLA-4 antibodies, including those used clinically, do not distinguish between the membrane-bound and soluble isoforms of CTLA-4 (sCTLA-4), and this limits the scope for unravelling the contribution of sCTLA-4 in functional and therapeutic studies.

Several hypotheses have been proposed to explain the mechanisms of anti-CTLA-4 immunotherapy in cancer. These include deletion of intratumoral T_reg_ cells (12), modulation of T-cell receptor (TCR)-CD28 interaction (13) and regulating naïve T-cell activation and differentiation (14). However, variation in clinical response is observed, and some studies have suggested that sCTLA-4 secreted by tumour cells may be responsible. For example, elevated levels of sCTLA-4 is reported found in serum from patients with malignant melanoma (15,16), mesothelioma (17) and acute B lymphoblastic leukaemia (18). A retrospective analysis of a small cohort of metastatic melanoma patients demonstrated that patients with higher levels of sCTLA-4 levels were more likely to respond to anti-CTLA-4 antibody ipilimumab than those with lower levels (19). Although such correlative studies provide support for the clinical relevance of sCTLA-4 in cancer, the functional properties and feasibility of isoform-specific antibody targeting of this molecule for cancer immunotherapy remain to be fully explored.

Following generation and characterisation of selective anti-sCTLA-4 monoclonal antibodies raised against the unique C-terminal epitope of sCTLA-4, we have shown that it regulates certain cell-extrinsic aspects of CTLA-4 function associated with distal control of T-cell effector responses in both health and disease (20–23). Contrary to previous assumptions, sCTLA-4 is produced as part of the natural immune response and should therefore be considered an important candidate regulatory mediator (15,24). Here, for the first time, we provide functional evidence of the strong immunosuppressive activity of sCTLA-4 *in vitro* and *in vivo*, and demonstrate it can be effectively targeted by an isoform-specific antibody to elicit anti-tumour activity.

## Results and Discussion

### sCTLA-4 constrains T-cell activation *in vitro*

Gene variants (**Fig. 1a**) of CTLA-4 have been reported to have utility as predictive biomarkers for anti-CTLA-4 therapy, improved long term survival, and precipitation for immune related adverse events (irAEs) (15,19). These correlative studies were largely based on circulating serum levels of sCTLA-4, but the relevance and potential impact of sCTLA-4 mRNA expression in tumour tissue has not been investigated. Interrogation of publicly available bulk RNAseq datasets available from The Cancer Genome Atlas (27) revealed relative expression of membrane-bound versus soluble CTLA-4 within lung adenocarcinoma (LUAD) and skin cutaneous melanoma (SKCM) (**Fig S1**). In this analysis we found that the membrane-bound isoform was the more abundant RNA species in both tumour types, and there was a positive correlation in the expression levels of each isoform that was greater in SKCM than in LUAD (**Fig S1**). Thus, tumour cells from patients express sCTLA-4 mRNA where the produced protein may modulate anti-tumour immune responses. To understand the functional capacity of sCTLA-4 to modulate anti-tumour T-cell responses, we constructed expression vectors to generate tumour cells that constitutively secrete recombinant sCTLA-4 and created stable cell lines (**Fig. 1b-c**). sCTLA-4 has been predominantly reported as a monomeric entity due to the splicing of exons 2 and 4, and subsequent loss of a membrane-proximal cysteine residue at position 157 (11,28). This cysteine residue, present in the membrane-bound CTLA-4 is presumed crucial for homodimerisation, stable interaction with B7 ligands and thus potent immunosuppressive properties (29,30). However, the skipping of exon 3 in sCTLA-4 results in a reading frame shift, encoding an alternative cysteine residue which may also permit sCTLA-4 dimerisation (**Fig. 1d**) (11). Indeed, we found a higher molecular weight component, consistent with the formation of a disulphide bridge between the de-novo encoded cysteine residues resulting from the alternative splicing that generates sCTLA-4, indicative of dimeric sCTLA-4, in cell culture supernatants from HeLa-sCTLA-4 cells, which under reducing conditions disassociated to form a monomer with the expected ~20 kDa mass (**Fig. 1e**).

**Figure 1.**
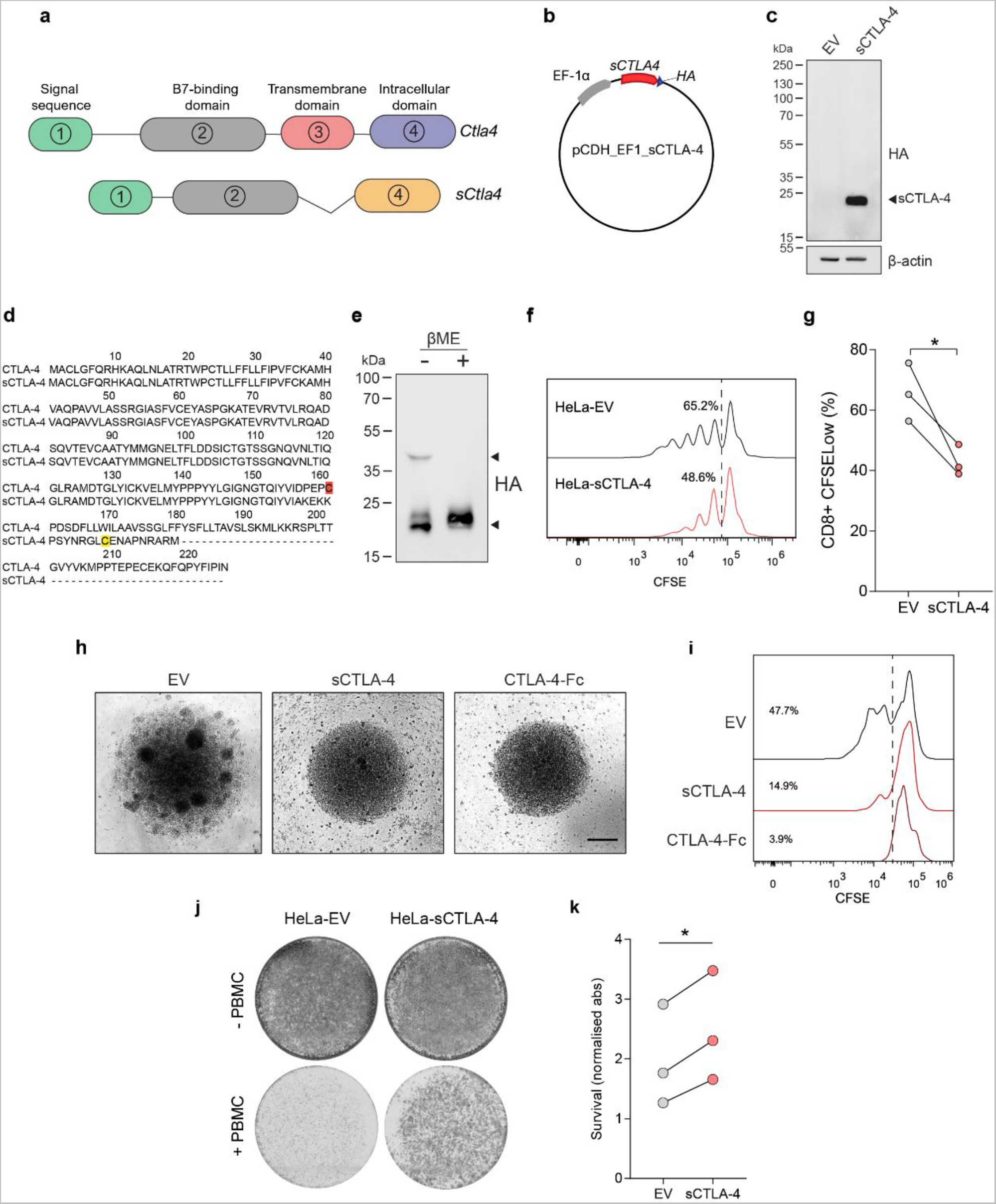
sCTLA-4 suppresses T cell activation and tumour cell killing *in vitro*. **a** Alternative splicing of CTLA-4 gives rise to soluble form of CTLA-4. Soluble CTLA-4 (sCTLA-4) retains the B7 binding domain encoded by exon 2 of membrane CTLA-4 but is missing the transmembrane domain encoded by exon 3. A reading frame shift during splicing gives rise to an alternative amino acid sequence encoded by exon 4. Both isoforms contain exon 1 which encodes a leader peptide. **b** Stable over-expression of recombinant sCTLA-4 in HeLa cervical adenocarcinoma cells. **c** Immunoblot showing transfected HeLa cells [HeLa-sCTLA-4 versus empty vector (EV)] under reducing conditions **d** Alignment of CTLA-4 and sCTLA-4 coding sequences. The membrane-proximal cysteine residue present in the transmembrane domain of membrane-bound CTLA-4 is highlighted in red. Although the transmembrane domain is absent in sCTLA-4, the new C-terminus in sCTLA-4 encodes another cysteine residue, highlighted in yellow. **e** Supernatant from HeLa-sCTLA-4 cells were immunoblotted under non-reducing and reducing conditions (−/+ βME). **f-g** Flow cytometric analysis of CFSE-stained PBMCs stimulated with anti-CD3 following co-culture with HeLa-sCTLA-4 or EV HeLa cells. Histograms indicate CD8^+^ T-cell proliferation after 4 days of anti-CD3 stimulation. Data represent 3 independent PBMC donors (* p<0.05 Student’s t-test). **h** Splenocyte-BMDM co-cultures following stimulation with PMA/ionomycin and treatment with recombinant CTLA-4-Fc or sCTLA-4-conditioned medium (scale = 50µm), representative of n=3. **i** Flow cytometric analysis of CD8^+^ T-cells from (h), histograms indicate CD8^+^ T-cell proliferation after 4 days. **j-k** T cell-mediated tumour cell killing assay of HeLa-sCTLA-4 cells. Images show crystal violet-stained viable HeLa cells following co-culture with anti-CD3 activated PBMCs. Data represent 3 independent PBMC donors (*p<0.05 Student’s t-test).

We next tested the immunosuppressive potential of sCTLA-4 on T-cell activation. HeLa-EV (empty vector control) or HeLa-sCTLA-4 cells were co-cultured with healthy donor human peripheral blood mononuclear cells (PBMCs) stimulated with anti-CD3 antibody. In this system, CD8^+^ T-cells exhibited reduced proliferation in the presence of sCTLA-4 secreting HeLa cells compared to those cultured with HeLa-EV control cells (**Fig. 1f-g**). A similar assay was performed using murine splenocytes treated with murine sCTLA-4 enriched supernatant. As a positive control within this assay we included a murine equivalent of Abatacept^TM^, which is a soluble recombinant CTLA-4-Fc fusion protein. Splenocytes stimulated with PMA/Ionomycin displayed clear clusters of proliferating cells that were suppressed when they were co-cultured either with murine sCTLA-4 or with CTLA-4-Fc (**Fig. 1h**). Furthermore, in keeping with our observations of human T-cell responses, specific examination of murine splenic CD8^+^ T-cells showed that the presence of murine sCTLA-4 or CTLA4-Fc reduced their proliferation compared to control cells (**Fig. 1i**). Taken together, these results show that sCTLA-4 is functionally similar to the artificially engineered CTLA4-Fc and suppresses T-cell immune responses.

Another aspect we investigated is whether sCTLA-4-induced constraints on T-cell activation translated into a survival advantage for tumour cells *in vitro*. Accordingly, HeLa cells were stained and visualised post-co-culture with activated PBMCs, revealing a markedly higher density of HeLa-sCTLA-4 cells than HeLa-EV cells, thereby indicating an enhanced resistance of target tumour cells to T-cell killing in the presence of sCTLA-4 (**Fig. 1j-k**). These data, taken together with our determined functional role of sCTLA-4, suggest that tumour release of sCTLA-4 gives malignant cells a survival advantage by suppressing immune cell responses. Moreover, since sCTLA-4 is evolutionarily conserved in mammals (11), the results from this section further suggest that human response to sCTLA-4 can be accurately modelled in mice.

### sCTLA-4 promotes syngeneic tumour growth *in vivo*

To confirm the immunosuppressive effects of sCTLA-4 in an immune-competent host, we monitored growth of murine syngeneic tumour cells constitutively expressing sCTLA-4 *in vivo*. We used B16F10 cells as a model of melanoma, creating cell lines expressing sCTLA-4 (B16F10-sCTLA-4) or empty vector (B16F10-EV). Both cell lines exhibited identical growth profiles *in vitro* (**Fig. 2a-b**), but when they were transplanted into mice B16F10-sCTLA-4 cells showed significantly accelerated growth compared to control (**Fig. 2c**). Similar results were observed in an experimental metastasis model where B16F10-sCTLA-4 cells spread into the lungs more efficiently and with higher burden than B16F10-EV control cells (**Fig. 2d**). In an alternative fibrosarcoma model where MCA-205 cells were modified to express sCTLA-4 (MCA-205-sCTLA-4), accelerated malignant cell growth resulting in an approximate 2-fold higher final tumour burden was observed in the mice transplanted with MCA-205-sCTLA-4 cells (**Fig. 2e-f**). Importantly, transplantation of either MCA-205-sCTLA-4 of MCA-205-EV cells into severely immunocompromised NOD SCID gamma (NSG) mice revealed overlapping growth profiles for the resulting tumours (**Fig. 2g-h**), indicating that the fitness advantage of MCA-205-sCTLA-4 tumours in syngeneic mice is not intrinsic to the cancer cells, but is rather dependent on extrinsic interaction with an intact immune mechanism.

**Figure 2.**
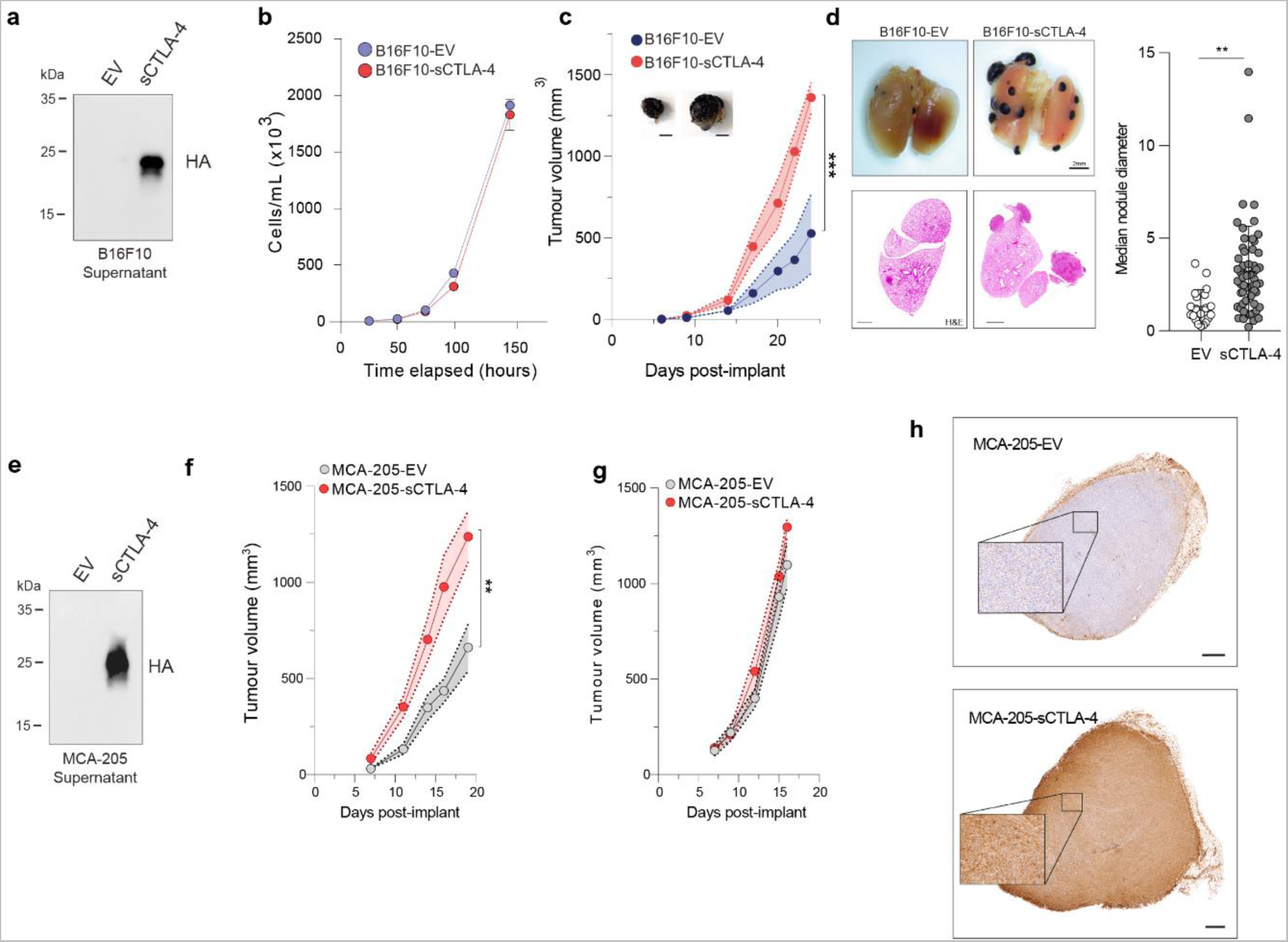
sCTLA-4 promotes syngeneic tumour growth *in vivo.* **a** Immunoblot showing stable sCTLA-4 over-expression in B16F10 melanoma cells. *In vitro* **(b)** and *in vivo* **(c)** growth curves of B16-EV versus B16-sCTLA-4 cells. *In vivo* tumour growth data are mean ± SEM, n=6 mice per group (p<0.001, Two-Way ANOVA). Representative photograph of tumours in each group are shown (scale = 50mm, n=6). **d** Experimental lung metastasis of mice inoculated intravenously with B16-EV or B16-sCTLA-4 tumours. Lung tumour nodule frequency and size was measured using stereomiscroscopy followed by H&E staining on day 26, with representative samples shown (scale = 1mm). Quantification data are mean ± SD, n=6 mice per group (p<0.01 two-tailed Student’s *t-*Test). **e** Immunoblot showing sCTLA-4 in cell culture supernatant from MCA-205-EV versus MCA-205-sCTLA-4 cells. **f** *In vivo* growth of MCA-205 tumours in immunocompetent and **g** immunocompromised NSG mice, n=7 mice per group. **h** Representative immunohistochemistry staining of MCA-205 tumours from **f** with anti-HA, scale bar = 0.5mm. Data are representative of at least two independent experiments.

### sCTLA4 raises the threshold for tumour infiltrating T-cell activation

The accelerated growth kinetics of sCTLA-4-secreting tumours was further investigated within the MCA-205 fibrosarcoma model. Here we focused on profiling the immune cells which infiltrate tumours using mass cytometry, comparing between mice exposed to MCA-205-sCTLA-4 and MCA-205-EV cells twenty days after implantation (**Fig. 3a**). Unsupervised clustering using flow self-organising maps (FlowSOM) was performed on data derived from live CD45^+^ cells within tumour infiltrates. This approach identified 14 cell populations of which 10 had characteristics that allowed assignment to distinct immune cell subsets, while 4 had characteristics which, although myeloid in nature, remained undetermined based on current understanding of myeloid cell phenotypes (**Fig. 3b**). These clusters were then visualised within t-distributed stochastic neighbour embedding (t-SNE) projections of the data to assess relative proportions of each subpopulation of cells (**Fig. 3c-d**).

**Figure 3.**
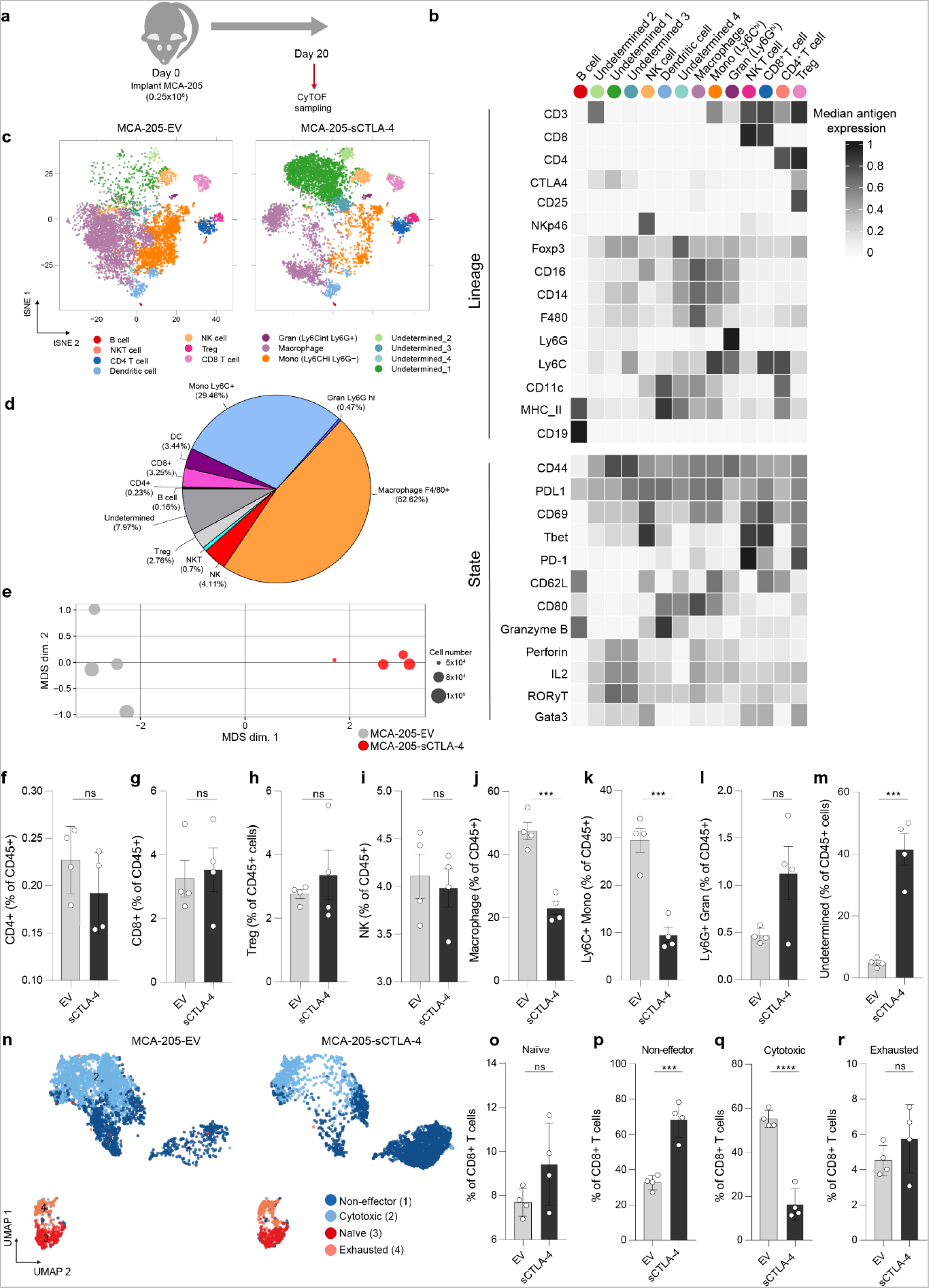
sCTLA-4 inhibits intratumoural T cell activation and differentiation. **a** Experimental design for mass-cytometric profiling of MCA-205 tumours day 20 post-inoculation. **b** Heatmap showing the median marker intensity of the 15 lineage markers used for flowSOM clustering of tumour infiltrates, in addition to functional state marker expression in each cluster. **c** tSNE analysis of the MCA-205 infiltrates. 25 flowSOM-identified metaclusters were manually merged according to lineage marker expression. Cells were proportionally combined from EV and sCTLA-4 expressing MCA-205 tumours (n=7 mice per group) to create the tSNE plot (1,000 cells per plot for visualisation). **d** The relative proportion of each flowSOM-derived metacluster. **e** PCA of MCA-205 tumours, dot size corresponds to cell number. **f-m** Comparison of the proportion of the indicated cell populations within tumour infiltrates. **n** UMAP of CD8^+^ T-cell subsets. CD8^+^ T-cells identified in (**b**) were re-clustered on functional state marker expression. The four clusters identified by flowSOM were manually annotated as naïve (CD62L^hi^CD44^−^), non-effector (CD44^+^ with low expression of CD69, IL-2, Perforin and Granzyme-B), cytotoxic (CD44^+^CD69^+^ expressing IL-2, Perforin and Granzyme B), and exhausted (PD-1^hi^) phenotypes. **o-r** Quantification of CD8^+^ T-cell subset frequency. Data in **f-m** and **o-r** are expressed as mean ± SD; *n =* 4 mice per group. Statistical significance was calculated using two-tailed Student’s *t*-test. Data is representative of two independent experiments.

Overall, myeloid subpopulations such as monocytes, dendritic cells (DCs) and macrophages represented the majority of the immune component of the tumour microenvironment (~30% and ~60% of CD45^+^ cells were monocytes and macrophages, respectively). Importantly, t-SNE projection of the data showed distinct changes to each subpopulation of cells associated with cell infiltrates from MCA-205-sCTLA-4 and MCA-205-EV tumours (**Fig 3c**), an observation that was confirmed by Principal component analysis (PCA) of median antigen expression values associated with all CD45^+^ cells from each model (**Fig. 3e**). Comparison of the immune cell composition within MCA-205-sCTLA-4 and MCA-205-EV tumours (**Fig. 3f-m**) revealed a dramatic reorganisation of the myeloid compartment, with reduced frequencies of both F4/80^+^ macrophages and Ly6C^hi^ monocytes in MCA-205-sCTLA-4 tumours (**Fig. 3j,k**). In contrast, within the overall lymphoid compartment, there were no differences in frequency of B and NK cells, or of CD4^+^, CD8^+^ T-cells and T_reg_ cells between MCA-205-sCTLA-4 tumours and EV controls (**Fig. 3f-i**). Importantly, the ratio of CD8^+^ to T_reg_ showed no change as well (**Fig. S2a**). The reduction on macrophages and monocytes occurred alongside a significant enrichment of an undetermined population of CD45^+^ cells characterised by expression of F4/80 and FoxP3 within infiltrates from MCA-205-sCTLA-4 tumours (**Fig. 3m**). This undetermined population of CD45^+^ cells retains myeloid features and is similar in phenotype to a subpopulation of F4/80^+^ / FoxP3^+^ macrophages reported to infiltrate lesions resulting from ischemic stroke (31). This report shows that such FoxP3^+^ macrophages have enhanced ability to scavenge debris, and so may have a similar role within the generated tumours of our model.

Our previous *in vitro* experiments showed that sCTLA-4 suppresses T cell activation, and we next investigated this within the population of tumour infiltrating T lymphocytes (TILs) in our system. To do this we examined CD8^+^ T-cell phenotypes by re-clustering according to functional state marker expression (32) (**Fig. 3n, Fig. S2b-c**). Notably, we observed a significant reduction in the percentage of cytotoxic CD8^+^ T-cells bearing potential effector activity in terms of granzyme B, perforin, and IL-2 and activation markers CD44^+^ and CD69^+^ in tumours from mice engrafted with MCA-205-sCTLA-4 tumours (**Fig. 3o-r**). In keeping with the reduction of cytotoxic CD8^+^ T-cells, there was enrichment in this group of tumours of naïve and non-effector T-cells typically lacking cytotoxic potential, whilst the proportion of exhausted PD-1^hi^ CD8^+^ T-cells was consistent between tumours expressing or lacking sCTLA-4. These results therefore confirm our in vitro data and show that secretion of sCTLA-4 by tumour cells actively suppresses T-cell activation in vivo. Collectively, these data suggest that sCTLA-4 profoundly modifies the immune context of the tumour microenvironment (TME) and dampens T-cell responses by raising the threshold for T-cell activation and attenuating T-cell effector function.

### Isoform-specific sCTLA-4 antibody augments anti-tumour immune responses

Having established that the presence of sCTLA-4 in the TME suppresses intratumoural T-cell activation and effector functionality *in vivo*, we sought to determine the effect of antibody-mediated sCTLA-4 blockade on immune cell profiles and T-cell responses within a transplantable syngeneic tumour. Genetic ablation of CTLA-4 in mice has been shown to result in multi-organ toxicity and lethal phenotype (33,34), whilst administration of naïve mice with pan-CTLA-4 antibody can lead to spontaneous development of autoimmune diseases (35). To ensure sCTLA-4 blockade did not induce such gross immune defects in normal immunocompetent hosts, we treated naïve C57BL/6 mice with anti-sCTLA-4 antibody twice a week for 5 weeks and assessed immune cell architecture in the primary and secondary lymphoid organs (thymus, spleen and lymph node) (**Fig. S3**). We detected no changes in the immune cell composition of these organs, providing assurance that administration of isoform-specific antibody does not disrupt overall immune homeostasis in naïve mice. We then treated mice transplanted with syngeneic MC38 colorectal tumours with anti-sCTLA-4 or Isotype control antibody every three days until day 21 (**Fig. 4a**). Anti-sCTLA-4 antibody treatment significantly attenuated tumour growth rate, inhibiting tumour growth by approximately 50% relative to control treated mice. To elucidate the cellular changes underlying this anti-tumour efficacy, tumour samples were collected for mass cytometry analysis at day 28 post-inoculation. Myeloid cell populations, including F4/80^+^ macrophages and Ly6C^hi^ monocytes, represented the major component of MC38 immune infiltrates (~85%) (**Fig. 4c-d and Figure S4a**), and were largely unaffected by treatment with anti-sCTLA-4 (**Fig 4i-l**). With respect to lymphoid cells, infiltrates of anti-sCTLA-4-treated tumours exhibited comparable frequencies of CD8^+^, CD4^+^ T-cells, T_reg_ cells and NK cells compared to control-treated tumours (**Fig. 4e-h**), with conservation of T-cell to T_reg_ ratios between the treatment groups (**Fig. S4b**). B lymphocytes seemed less abundant in anti-sCTLA-4 treated tumours relative to controls (**Fig 4m**), and there was clear reduction in an undetermined population of CD45^+^ cells characterised by F4/80 and FoxP3 expression within the infiltrates from anti-sCTLA-4 treated mice (**Fig. 4n**). To assess the effect of sCTLA-4 inhibition on T-cell function, CD8^+^ T-cells were re-clustered according to state marker expression (**Fig. 4o and Fig. S4c-d**). We observed fewer CD8^+^ T-cells designated as having non-effector function but an enrichment of activated CD69^+^ effector T-cells expressing IL-2, granzyme B and perforin in anti-sCTLA-4 treated tumours, indicating that sCTLA-4 blockade promotes the activation and differentiation of TILs (**Fig. 4o-r**). Taken together with the results of the previous section, these data suggest that anti-sCTLA-4 antibodies can reverse the effects of sCTLA-4 and restore immune response to tumours.

**Figure 4.**
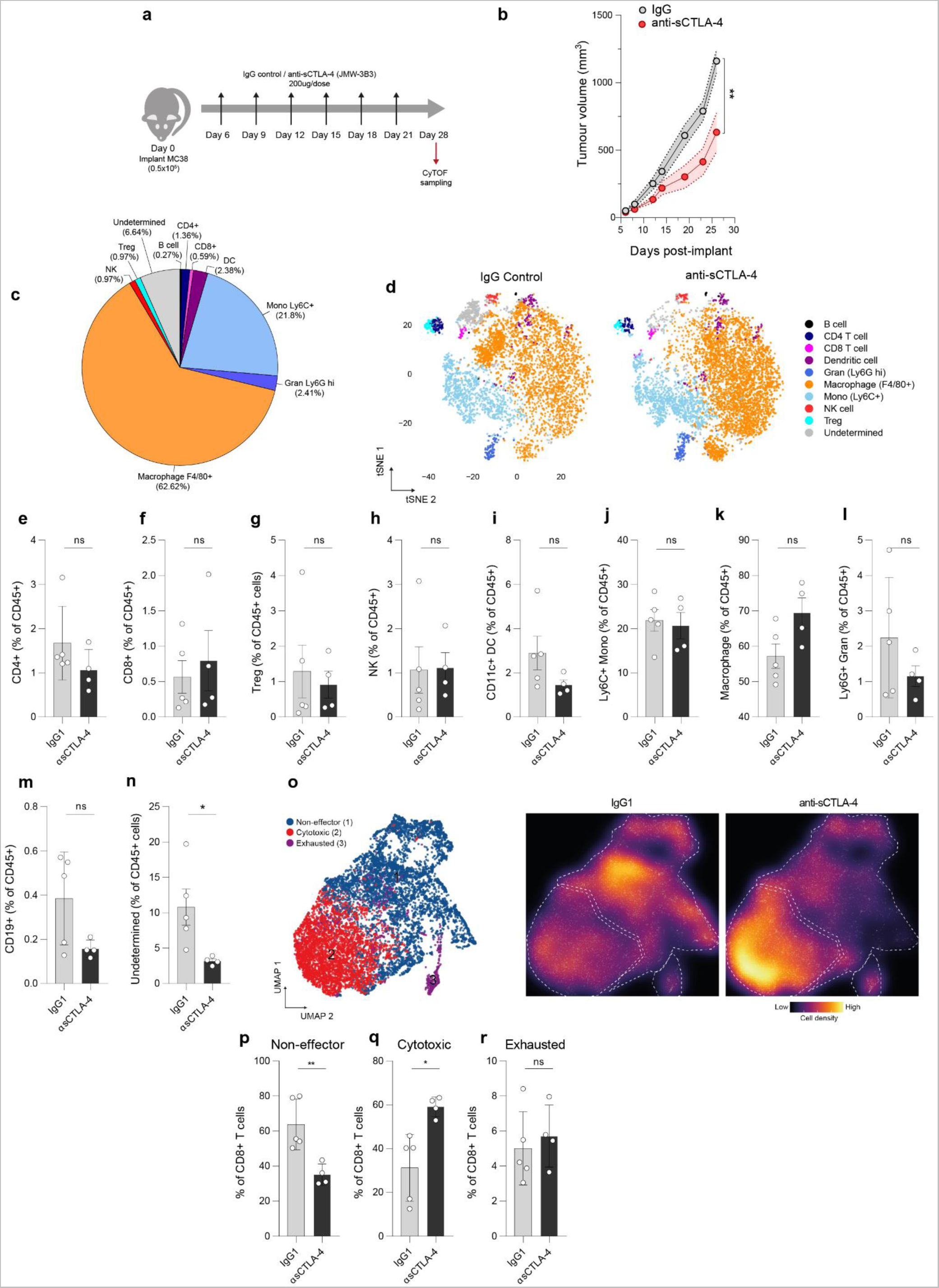
sCTLA-4 blockade promotes T-cell cytotoxic function and attenuates murine tumour growth *in vivo.* **a** Schematic showing study design and schedule for the treatment of mice bearing MC38 tumours. **b** *In vivo* growth of tumours treated according to **(a)**. Data are mean ± SEM, n=16-18 mice per group, from 3 independent experiments (** p<0.01, *** p<0.001, Two-Way ANOVA). **c** Proportions of major tumour-infiltrating leukocyte populations, expressed as percentage of CD45^+^ cells. **d** tSNE analysis of immune infiltrates isolated from MC38 tumours in (**b**) at day 28. Cell lineage markers were used for flowSOM-based meta-clustering of CD45^+^ cells. 25 flowSOM-identified metaclusters were manually merged according to lineage marker expression. Clustering was performed on all cells from both treatment groups (n=4-6 mice per group), with 1,000 cells visualised in tSNE plots. **e-n** Comparison of proportions of the indicated cell populations between anti-sCTLA-4 or isotype control treated tumours. **o** UMAP of CD8^+^ T-cell subsets. CD8^+^ cells identified in **d** were re-clustered using functional state marker expression. Three clusters identified by flowSOM were manually annotated as non-effector (CD44^+^ with low expression of CD69, IL-2, Perforin and Granzyme-B), cytotoxic (CD44^+^IL-2^+^Perforin^+^Granzyme-B^+^), and exhausted (PD-1^hi^) phenotypes. Shifts in CD8 functionality following treatment, visualized by cell density scale on the CD8^+^ T-cell subset UMAP. **p-r** Quantification of CD8^+^ T-cell subset frequency in treated tumours. Data in **e-n** and **p-r** are expressed as mean ± SD; n=4-6 in each group. Statistical significance was calculated using two-tailed Student’s *t*-test. Representative of 2 independent experiments.

Despite growing clinical applications of anti-CTLA-4 antibodies for the treatment of cancer, its precise mechanisms of action remain poorly defined. The current perception of CTLA-4 biology focuses almost entirely on the membrane-bound receptor, and it is the activity of this isoform that almost all studies focus on to elicit greater anti-tumour immune responses. As a result, there has been a tendency to assume that checkpoint regulation operates solely through membrane-based interactions, dismissing any contribution of the soluble isoform in this process. However, our data provide direct evidence that sCTLA-4 mediates modulation of murine and human T-cell activation in a range of situations, exhibiting function similar to that of the engineered soluble dimeric fusion protein CTLA4-Fc. We show that transforming cancer cells to secrete sCTLA-4 in the TME promotes tumour growth and metastasis by limiting CD8^+^ T-cell effector activity, demonstrating the functional significance of this less characterised isoform. Some cancer cells have been found to produce sCTLA-4 naturally (15) (36) (37) and we predict that this may lead to blunted T-cell effector activity and immune escape. This is further supported by the finding that cancer cell-intrinsic expression of sCTLA-4 did not provide growth advantage to cancer cells in vitro, or in an immunocompromised host - evidence that largely favours the hypothesis that immune cells, particularly CD8^+^ T-cells are being held at bay by sCTLA-4. Anti-sCTLA-4 treatment enriched the proportion of tumour-infiltrating CD8^+^ T-cells with cytotoxic functionality, likely by overcoming the higher threshold exerted by sCTLA-4 for CD8^+^ T-cell activation and effector function. Given the immunological effects achieved by blocking sCTLA-4 and that conventional anti-CTLA-4 antibodies such as ipilumimab concurrently block both membrane-bound and sCTLA-4, it is likely that their anti-tumour activity occurs at least in part through blockade of sCTLA-4. Acknowledging the relative role of sCTLA-4 in health and disease may lead to distinct interpretation of many observations concerning membrane-bound CTLA-4 function, relevant across an enormous range of clinical applications including cancer, autoimmunity, allergy and transplant biology.

## Materials and Methods

### Cell lines

Mouse tumour cell lines MC38 and B16F10 were obtained from the ATCC (London, UK) whilst MCA-205 were from Sigma-Aldrich. Human HEK 293T cells were obtained from ATCC. Cells were cultured in RPMI-1640 or DMEM containing 10% foetal-bovine serum (FBS) and 1% penicillin/streptomycin (Gibco). All cell lines were routinely tested for mycoplasma infection by PCR. Human peripheral blood mononuclear cells (PBMC) were purified from healthy donor leukocyte cones purchased from Blood and Transplant, Speke, Liverpool by Lymphoprep™ (Serumwerk) density centrifugation and cultured in RPMI-1640 + 10% FCS/1% penicillin/streptomycin.

### Mice

All animal studies were performed under UK Home Office Project Licence PP6634992, in accordance with the UK Animal (Scientific Procedures) Act 1986 and the EU Directive 86/809. Studies were approved by the University of Liverpool Animal Welfare and Ethical Review Body (AWERB) and referred to the Workman guidelines (25).

### Preclinical tumour models

Tumour cells were harvested in log-phase growth prior to detachment with 0.25% Trypsin-EDTA (Gibco) solution and washing plus resuspension in PBS (Gibco). For *in vivo* growth studies, 5×10^5^ B16F10 or MC38 or 2×10^5^ MCA-205 were injected subcutaneously into the right-flank of female C57BL/6J or NOD-SCID gamma (NSG) mice (6-8 weeks of age). Tumour growth was measured using callipers and the volume formula (width^2^ × length)/2, with a tumour burden limit of 1500 mm^3^. For experimental metastasis models measuring B16F10 lung engraftment, 2×10^5^ cells were injected into the tail-vein of 6-8 week-old female C57BL/6 mice.

### Immunoblot

Cell extracts were prepared using RIPA buffer (Thermo Fisher Scientific), or cell culture supernatant and diluted in sample buffer with β-mercaptoethanol. Information on antibodies used is given in supplementary table 1. After SDS-PAGE, proteins were transferred onto PVDF membranes (Merck) and blocked using 5% non-fat dried milk in PBS-Tween 20. Following overnight incubation in primary antibody, membranes were washed and incubated in HRP-conjugated secondary antibody before detection using Immobilon ECL Ultra substrate (Millipore).

### *In vitro* co-cultures

Human PBMCs were grown in RPMI-1640 medium with 10% FBS + 1% penicillin-streptomycin before staining with 5µM cell-tracker CFSE (Invitrogen). PBMCs and HeLa target cells were co-cultured in u-bottom 96 well plates at a range of T:E ratios (indicated in figure legends) T cells were stimulated with 10 µg/mL plate-immobilised anti-CD3 (OKT3, BioLegend) and cultured for 4 days prior to staining with anti-CD8-APC (SK1, BioLegend) and proliferation measured by flow cytometry, judged by CFSE dilution. For T-cell killing assays, following PBMC-HeLa co-cultures, suspension cells were removed by PBS washes. HeLa cells which remained viable and adherent were then fixed in absolute methanol (Sigma-Aldrich) for 20 minutes at room temperature, before staining with 0.5% crystal violet solution (Sigma-Aldrich).

### *In vitro* T cell suppression assay

To measure the suppression of T-cell proliferation by sCTLA-4, CFSE-stained splenocytes were co-cultured with BMDMs (20:1 splenocyte to BMDM) in RPMI-1640 + 10% FCS + 0.05mM β-mercaptoethanol. Cultures were stimulated with phorbol 12-myristate 13-acetate (PMA) (5 ng/mL, Alfa-Aesar) + Ionomycin (0.5 µg/mL, Thermo Fisher Scientific) and supplemented with sCTLA-4 conditioned media or murine CTLA-4-Fc fusion protein (10 µg/mL, BioLegend). After 4 days, cells were harvested and stained with anti-mouse-CD8α (BioLegend) for analysis of T cell proliferation by flow cytometry as above.

### BMDM isolation and culture

Femurs and tibias from 6–8-week-old female C57BL/6J mice were flushed with ice-cold PBS into petri-dishes. The suspension was then pressed through 70µm cell strainers (Corning) before centrifugation at 200xg for 5 minutes at 4°C. Bone marrow cells were differentiated into BMDM by culturing them into DMEM+10%FCS+25ng/mL recombinant murine M-CSF, BioLegend) to a density of 1×10^6^ cells/mL for 7 days, with the culture medium being replaced every 2 days.

### Cloning

Expression plasmids were constructed for the stable over-expression of human and mouse sCTLA-4 in various tumour cell lines. Human and murine sCTLA-4 open-reading frames (ORF) were PCR-amplified from the cDNA of anti-CD3 activated PBMCs or splenocytes, respectively. Amplification of ORFs used the following primer sequences: 5’-ATGGCTTGCCTTGGATTTCAG-3’ (human sCTLA-4 forward), 5’-AGTCACATTCTGGCTCTGTTGG-3’ (human sCTLA-4 reverse), 5’-ATGGCTTGTCTTGGACTCCG-3’ (murine sCTLA-4 forward) and 5’-TCACATTCTGGCTCTGTTGG-3’ (murine sCTLA-4 reverse). The sCTLA-4 ORF was cloned into a pCDH-EF1-FHC lentiviral expression vector by restriction digest and ligation with T4 DNA ligase (Thermo Fisher Scientific), to produce pCDH-EF1-sCTLA-4 expression vectors. pCDH-EF1-FHC was a gift from Richard Wood at MD Anderson Cancer Centre, University of Texas, US (Addgene plasmid #64874).

### Generation of stable expression cell lines

For lentivirus generation, 4×10^5^ HEK 293T cells were transfected with pCDH-EF1-sCTLA-4 expression plasmid (1.5 µg) and psPax2 (2 µg) and pMD2.G (1.5 µg) packaging plasmids (Addgene), using Viafect™ (Promega) and OptiMEM (Gibco). Media was replaced on the following morning with fresh DMEM+10% FCS. Lentiviral supernatant was collected 48-72 hours post-transfection, clarified by centrifugation for 500xg for 5and minutes and filtered using 0.45 µm PES filters (Starlab). For lentiviral transduction, 2×10^5^ target tumour cells were seeded in 6 well plates and infected with lentiviruses + polybrene (8 µg/mL) before selection of transduced cells using 4ug/mL puromycin (Sigma-Aldrich).

### Isolation of tumour-infiltrating leukocytes

Excised tumours were mechanically disrupted and incubated with 1.67 Wünsch U/mL Liberase TL (Roche) and 0.2 mg/mL DNase I (Merck) for 30 minutes at 37°C with agitation. Digested tumours were homogenised by pipetting before being passed through 100µm nylon mesh strainers (Corning). Cell suspensions were washed with RPMI-1640 before proceeding to mass cytometry staining.

### Mass cytometric immunophenotyping

Custom antibody-metal conjugations were prepared using the Maxpar Antibody Labeling Kit (Standard Biotools) according to the supplied protocol recommendations. Afterwards, Maxpar-conjugated antibodies were stored in PBS-based antibody stabilization solution (Candor Biosciences) at 4°C and titrated before use. Isolated TILs were washed with PBS prior to viability staining with cisplatin (Standard Biotools). Samples were then barcoded using metal-labelled anti-mouse-CD45, and an equal number of cells from each sample was pooled for subsequent staining. Samples were incubated with TruStain FcX™ (anti-mouse CD16/32) for 10 minutes on ice to block Fc receptors before immunostaining with antibodies for lineage and state-defining markers (given in supplementary table 2). Samples were then washed with cell staining buffer (Standard Biotools) before fixation in 4% paraformaldehye (Thermo Fisher Scientific). For measurement of intracellular cytokines, samples were processed using the FoxP3/Transcription factor fix-perm kit (eBioscience). Samples were incubated with Cell-ID-intercalator-Ir (Standard Biotools) for doublet discrimination prior to analysis. Data were pre-processed to isolate live CD45^+^ cells using FlowJo (BD) before analysis and unsupervised clustering was performed using the R-based package CATALYST (26).

### Immunohistochemistry

Tumours were excised and fixed in 10% neutral-buffered formalin for 48 hours at 4°C. Deparaffinization and antigen retrieval on sections was performed using the Dako PT-link station (Agilent), before immunostaining overnight at 4°C with anti-HA antibody (CST). For secondary staining, an HRP labelled polymer from the EnVision+ system was used (Dako). Staining was developed using diaminobenzidine and counterstained with haematoxylin (Sigma-Aldrich).

### RNA-seq patient datasets

Normalised isoform-level data RNA sequencing (RNA-seq) data were obtained through The Cancer Genome Atlas (TCGA) Data Matrix portal (Level 3, tcga-data.nci.nih.gov/tcga/dataAccessMatrix.htm) and Firebrowse (firebrowse.org/). Transcript isoform identification codes were mapped to gene names using the UCSC table browser (27).

### Statistical analysis

Data are expressed as means ± SEM unless otherwise indicated. Student’s t test and two-way ANOVA were performed for the statistical analysis and the details can be found in the figure legends. For comparison of tumour volumes, repeated measures two-way ANOVA followed by Sidak’s test (two groups) was used. Statistical analysis was performed with GraphPad Prism (ver. 9.5.0, GraphPad Software). *P* values less than 0.05 were considered significant.

### Data availability

All data generated present in the figures and supplementary material. Materials available upon reasonable request.

## Acknowledgements

The authors would like to thank Southampton Antibody and Vaccine Group (Christine Penfold, Kerry Cox and Martin Taylor) for the production and shipping of anti-sCTLA-4 antibody JMW-3B3.

## Figure legends

**Figure S1.**
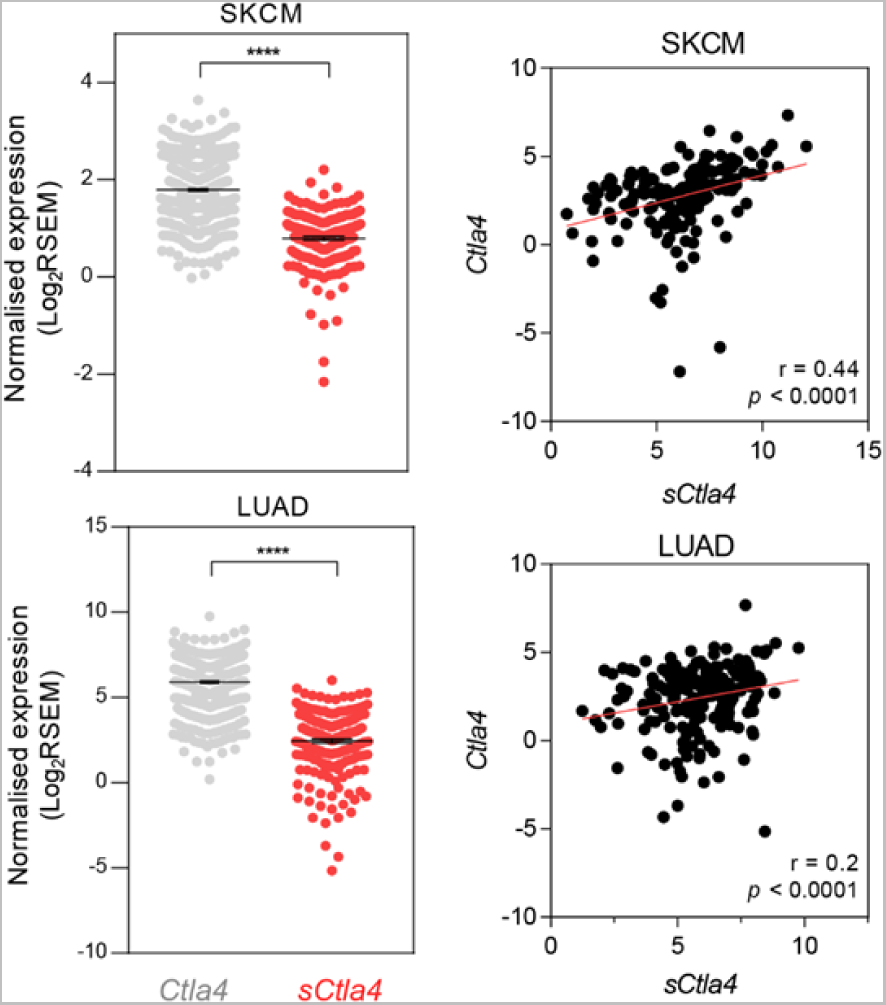
sCTLA-4 expression correlates with membrane CTLA-4 isoform levels. Quantification of CTLA4 isoform mRNA levels in SKCM and LUAD TCGA bulk tumour RNAseq datasets (**** p<0.0001 Kolmogorov-Smirnov test) and correlation analysis of bulk tumour soluble and membrane-bound CTLA4 mRNA. Pearson’s coefficient is given.

**Figure S2.**
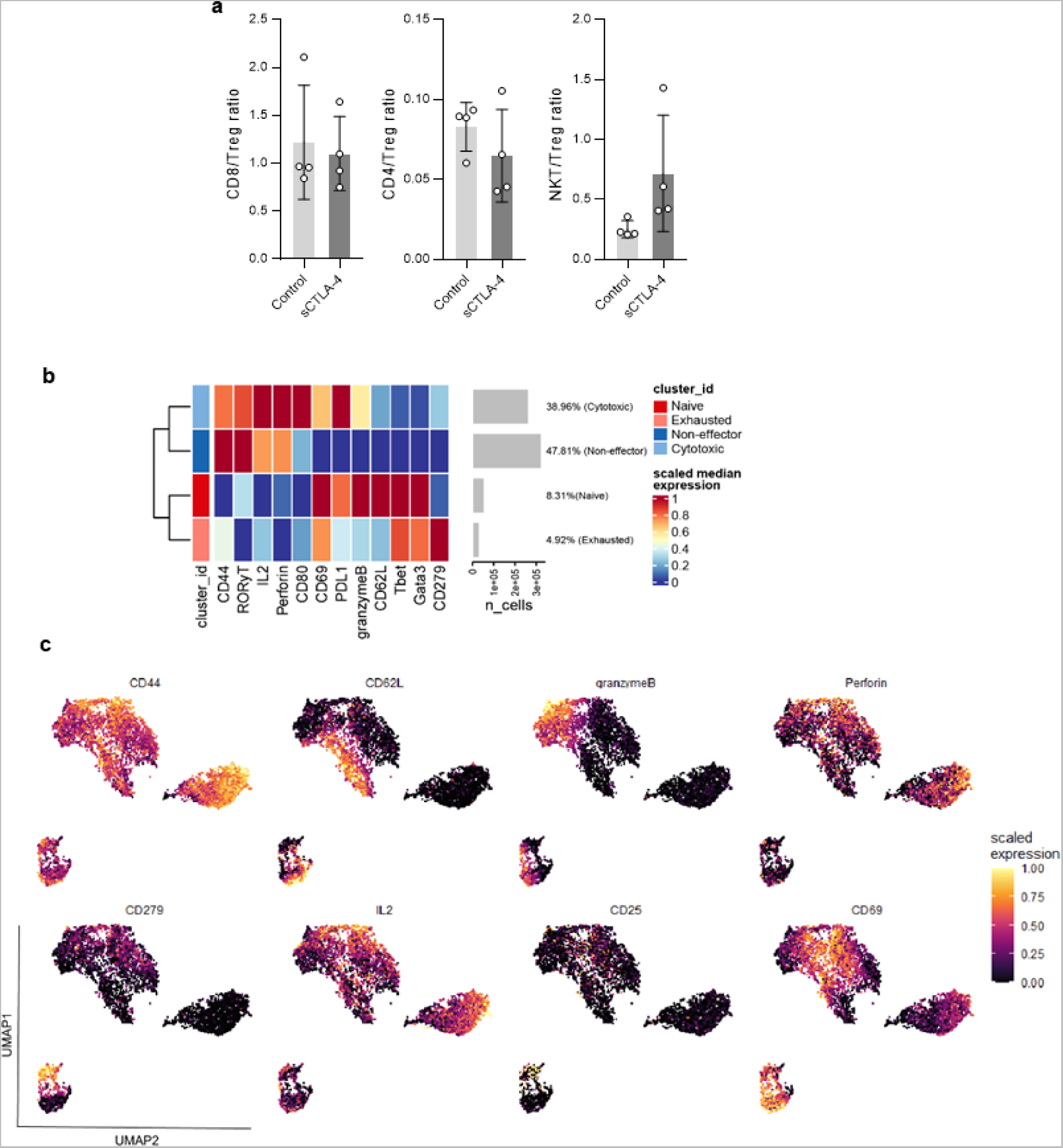
Lymphocyte:Treg ratios and scaled CD8 T-cell marker expression in MCA-205 tumours. **a** Ratio of CD8, CD4 and NKT to Treg in tumour infiltrates bearing MCA-205-EV control or MCA-205-sCTLA-4 tumours. Data are expressed as mean ± SD; n = 4 mice per arm. Statistical significance was calculated using two-tailed Student’s t-test. **b** Heatmap showing state marker expression in CD8^+^ T-cell clusters and **c** CD8^+^ T-cell UMAPs coloured by state marker expression.

**Figure S3.**
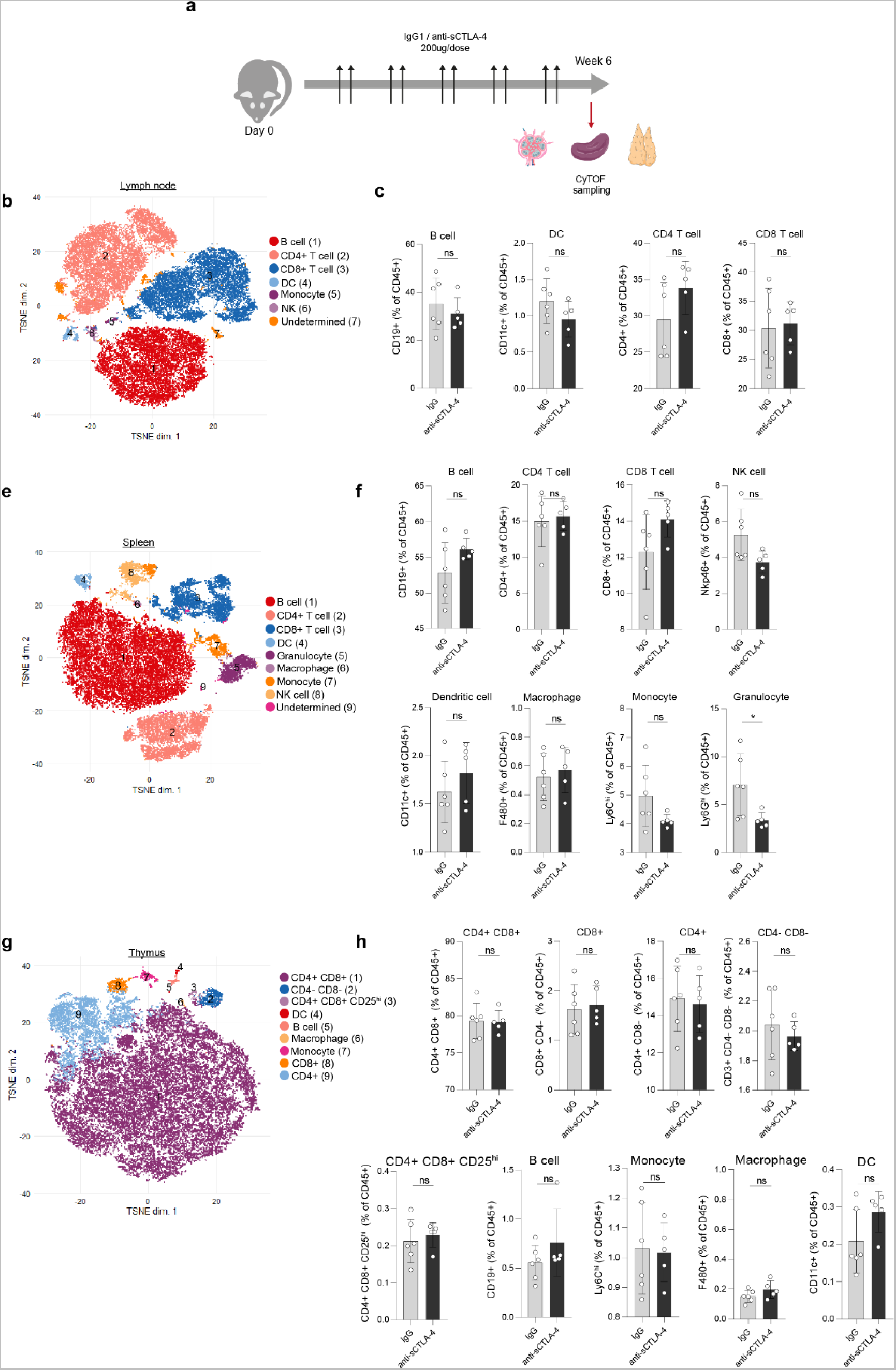
Anti-sCTLA-4 treatment does not perturb immune homeostasis. **a** Five-week-old C57Bl/6 mice were treated twice weekly for 5 weeks with 200ug of anti-sCTLA-4 (JMW-3B3) or Isotype control antibody. **b,e,g** Mass cytometric analysis of mice in a: tSNE analysis showing flowSOM based clustering of major cell populations within lymph node, spleen, and thymus respectively. **c,f,h** Quantification of major cell subsets within these organs. Data are based on aggregated scaled expression, n=4-6 mice. 2,000 cells per tSNE plot are displayed.

**Figure S4.**
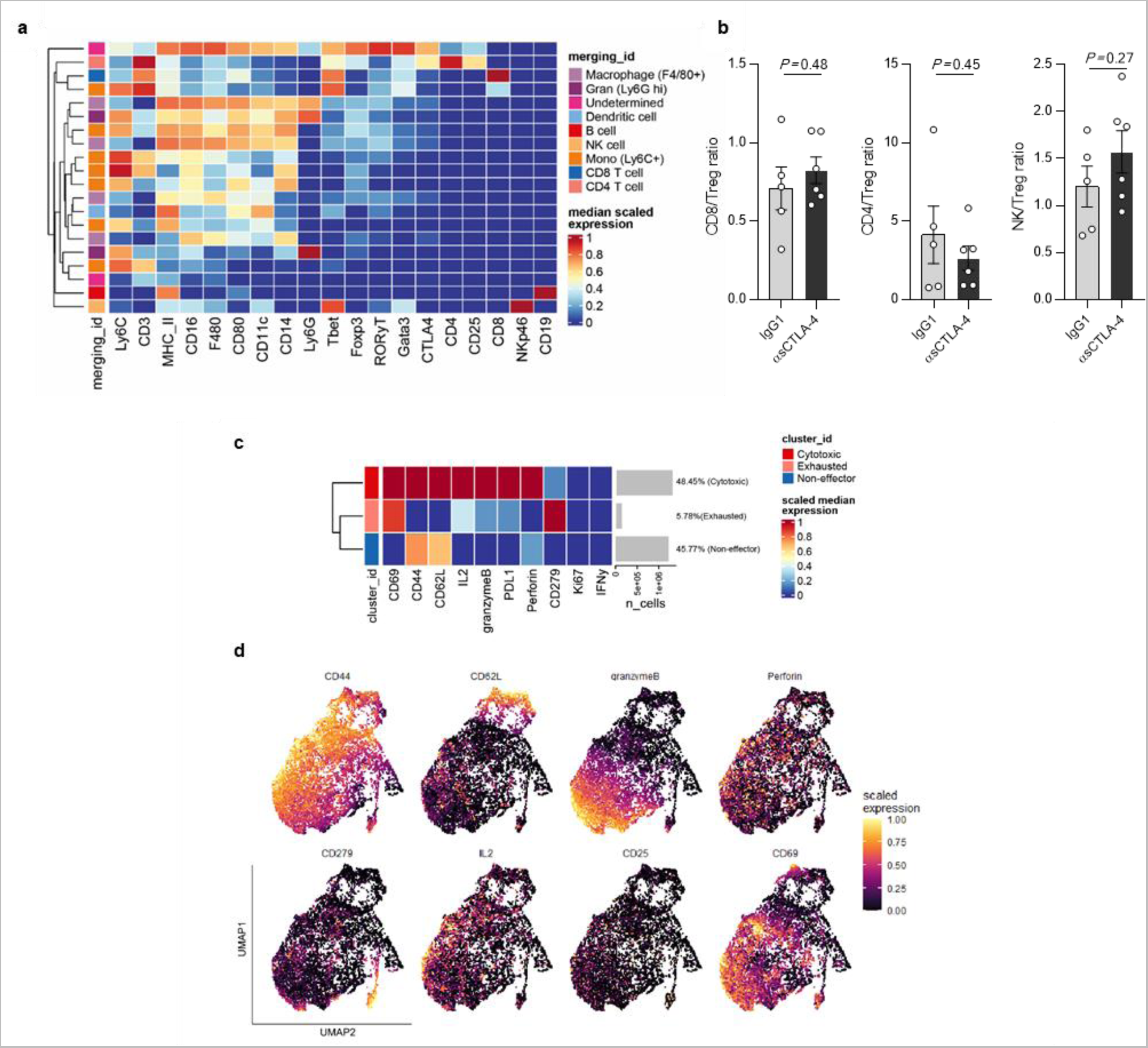
Evaluation of MC38 derived immune infiltrate clusters by mass cytometry. **a** Heatmap showing scaled marker expression in manually annotated clusters from data shown in Fig. 4c. 25 flowSOM-identified metaclusters were manually merged according to lineage marker expression. **b** Lymphocyte: T_reg_ cellratios of MC38 tumour bearing mice treated with anti-sCTLA-4 or isotype control antibody. Data are expressed as mean ± SD; n = 4 mice per arm. Statistical significance was calculated using two-tailed Student’s t-test. **c** Scaled CD8^+^ T-cell marker expression in MC38 tumours with heatmap showing state marker expression in CD8^+^ T-cell clusters. **d** CD8 T cell UMAPs coloured by state marker expression.

**Supplementary Table 1.** Antibodies and recombinant proteins used in the study.

| Antibody / protein | Clone | Supplier (cat #) | Usage |
| --- | --- | --- | --- |
| Anti-β-actin | AC-15 | Sigma-Aldrich (A1978) | Immunoblot |
| Anti-CTLA-4 | E1V6T | CST (96399) | Immunoblot |
| Anti-HA | C29F4 | CST (3724) | Immunoblot / IHC |
| Anti-rabbit IgG, HRP-linked |  | CST (7074) | Immunoblot |
| Anti-mouse IgG, HRP-linked |  | CST (7076) | Immunoblot |
| Anti-human-CD8-APC | SK1 | BioLegend (344722) | Flow cytometry |
| Anti-human-CD4-APC | SK3 | BioLegend (980802) | Flow cytometry |
| Anti-mouse-CD8a-APC | 53-6.7 | BioLegend (100711) | Flow cytometry |
| Anti-mouse-CD4-FITC | RM4-5 | BioLegend (100510) | Flow cytometry |
| CTLA-4-Fc |  | BioLegend (591802) | Functional study |
| Anti-sCTLA-4 | JMW-3B3 | Custom | Functional study |
| InVivoMAb anti-mouse CTLA-4 (CD152) | 9D9 | BioXcell (BE0164) | Functional study |
| InVivoMAb mouse IgG2b | MPC-11 | BioXcell (BE0086) | Functional study |
| InVivoMAb mouse IgG1 | MOPC-21 | BioXcell (BE0083) | Functional study |

**Supplementary Table 2.**
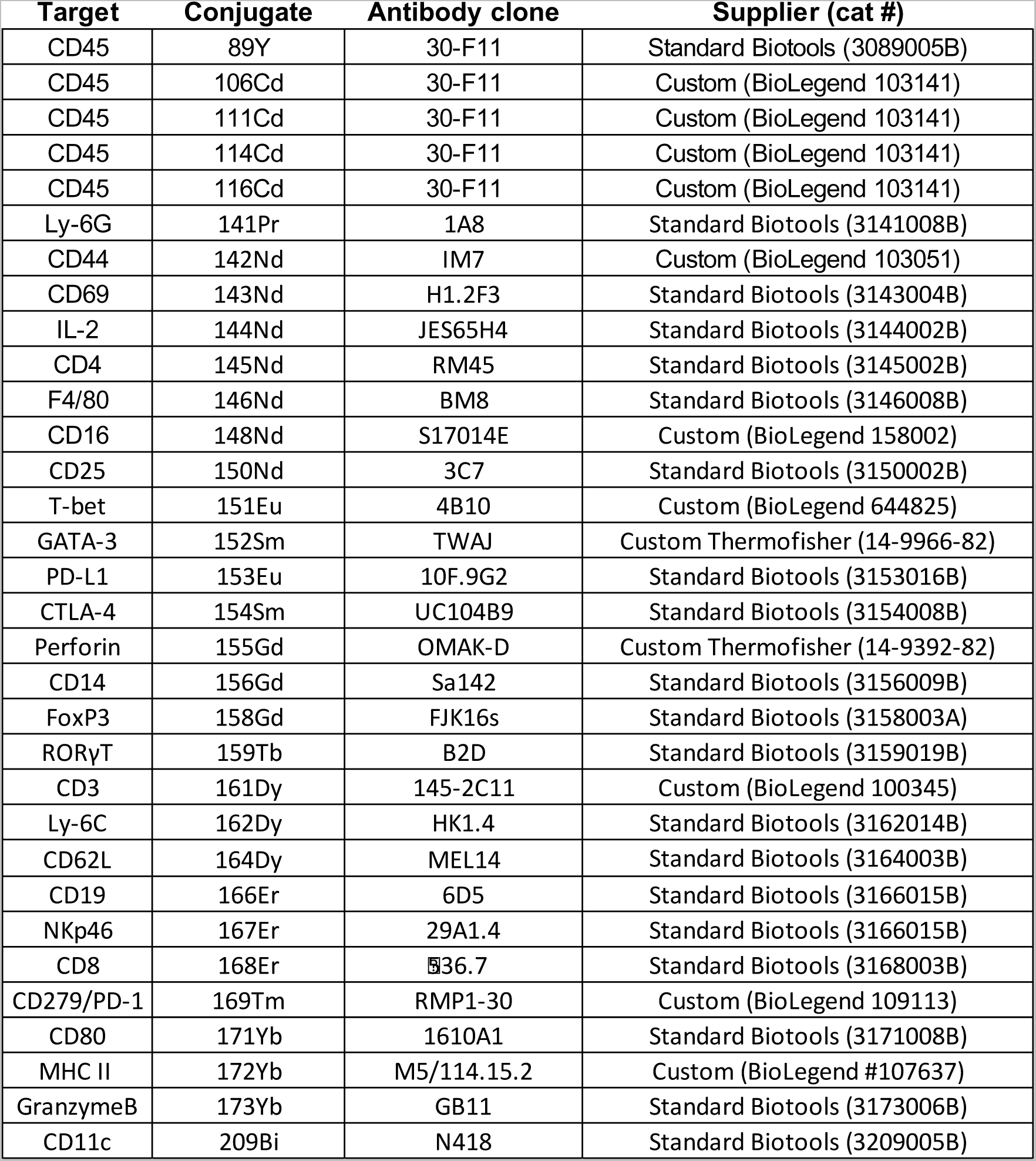
Mass cytometry antibodies for immunophenotyping.

